# The causal role of circulating vitamin D concentrations in human complex traits and diseases: a large-scale Mendelian randomization study

**DOI:** 10.1101/677922

**Authors:** Xia Jiang, Tian Ge, Chia-Yen Chen

## Abstract

Vitamin D has been associated with a variety of human complex traits and diseases in observational studies, but a causal relationship remains unclear. To examine a putative causal effect of vitamin D across phenotypic domains and disease categories, we conducted Mendelian randomization (MR) analyses using genetic instruments associated with circulating 25-hydroxyvitamin D [25(OH)D] concentrations. We leveraged genome-wide significant 25(OH)D-associated SNPs (N=138) from a meta-analysis combining a vitamin D GWAS conducted in 401,460 white British UK Biobank (UKBB) participants and an independent vitamin D GWAS including 42,274 samples of European ancestry, and examined 190 large-scale health-related GWAS spanning a broad spectrum of complex traits, diseases and biomarkers. We applied multiple MR methods to estimate the causal effect of vitamin D while testing and controlling for potential biases from horizontal pleiotropy. Consistent with previous findings, genetically predicted increased 25(OH)D levels significantly decreased the risk of multiple sclerosis (OR=0.824; 95%CI=0.689-0.986). The protective effect estimate was consistent across different MR methods and four different multiple sclerosis GWAS with varying sample sizes and genotyping platforms. On the contrary, we found limited evidence in support of a causal effect of 25(OH)D on anthropometric traits, obesity, cognitive function, sleep behavior, breast and prostate cancer, and autoimmune, cardiovascular, metabolic, neurological and psychiatric traits and diseases, and blood biomarkers. Our results may inform ongoing and future randomized clinical trials of vitamin D supplementation.

## Introduction

Vitamin D (or its hydroxylated form produced by liver, the 25-hydroxyvitamin D [25(OH)D]) is a group of fat-soluble secosteroids that pose a crucial impact on human health.^1^ Long-term poor dietary intake of vitamin D or inadequate exposure to sunlight (ultraviolet B) can cause rickets and osteomalacia. Several large-scale epidemiological investigations have linked vitamin D deficiency to a range of common diseases such as cancer, autoimmune diseases, and cardiovascular conditions.^2–6^ Yet the observational nature of epidemiological designs hinders causal inference, and the validity of these results can be plagued by measurement error, confounding and reverse causality. Although randomized controlled trials (RCTs) are considered as the gold standard to establish causal relationships, large-scale RCTs of vitamin D supplementation are not readily available due to their high cost, long duration and ethical concerns.^7,8^ A recently completed RCT, VITAL (ClinicalTrials.gov: NCT01169259), which involved a total of 25,871 participants with a median follow-up period of 5.3 years, did not find lowered incidence of invasive cancer or cardiovascular events for those who took vitamin D3 at a dose of 2,000 IU per day compared with the placebo group.^7,9^ The ancillary study to VITAL, VITAL-DEP (ClinicalTrials.gov: NCT02521012), recruited and followed up with 18,353 participants, and found the risk of depression or clinically relevant depressive symptoms was not significantly different between the vitamin D3 group and the placebo group; no significant benefit of vitamin D was identified for depression incidence or recurrence either.^10^ RCTs targeting other diseases and conditions such as cognitive change (ClinicalTrials.gov: NCT03613116) are ongoing and have yet to reach conclusions. To better inform the field on the chance of success for such vitamin D trials, an improved understanding of the causal role of vitamin D in human complex traits and diseases, preferably through less costly observational studies, is needed.

Serum levels of 25(OH)D has a polygenic predisposition. Several early genome-wide association studies (GWAS) have demonstrated the associations of *GC, NADSYN1/DHCR7, CYP2R1, CYP24A1, SEC23A,* and *AMDHD1* with circulating vitamin D.^11–14^ One of the latest GWAS of circulating 25(OH)D, conducted by meta-analyzing a GWAS of 401,460 white British UK Biobank (UKBB) participants and a GWAS of 42,274 samples of European ancestry, has identified 138 conditionally independent SNPs in 69 vitamin D associated loci.^13^ These genetic discoveries have enabled causal inference between vitamin D and complex traits through Mendelian randomization (MR), an approach that correlates genetic variants randomly distributed at conception (instrumental variables, IVs) with health outcomes later on.^15^ An unbiased estimate of the causal effect can be obtained using the observed IV-exposure and IV-outcome associations when all model assumptions of MR are satisfied.^16^

To date, the causal role of vitamin D beyond its importance in bone health remains ambiguous and is under much debate. The causal effects of vitamin D were only replicated in MR studies for adult- and pediatric-onset multiple sclerosis and Alzheimer’s disease;^17–21^ whereas findings for other complex human traits and diseases were inconsistent.^22–25^ Overall, results from MR studies of vitamin D are largely null and do not support conventional observational studies, which have associated vitamin D with a variety of human traits and diseases. That said, it is worth noting that the majority of hitherto MR studies for vitamin D used a small number of IVs (N=4 or 6 independent loci) identified by the SUNLIGHT consortium meta-GWAS back in 2010.^10^ Recent GWAS incorporating UKBB samples have identified a much larger number of 25(OH)D-associated genomic loci with more accurate effect size estimates, providing an opportunity to reevaluate the causal role of 25(OH)D with much improved power.^13,14^ Moreover, the rapid accumulation of GWAS summary statistics for human complex traits and diseases has allowed us to assess the causal effect of vitamin D across a broad spectrum of health-related outcomes.

Here, we conducted two-sample MR analyses (i.e., IV-exposure and IV-outcome associations are estimated from two separate samples) to examine the causal role of 25(OH)D across phenotypic domains and disease categories. Summary statistics for 138 independent genetic instruments were extracted from one of the latest and largest vitamin D meta-GWAS incorporating UKBB samples. Summary statistics for the IV-outcome associations were obtained from 190 large-scale GWAS spanning complex traits, diseases and biomarkers.

## Methods

### Data for IV-exposure

We extracted summary statistics for SNPs significantly associated with circulating 25(OH)D concentration from a meta-analysis of two vitamin D GWAS conducted by Manousaki et al.: a GWAS in UK Biobank including 401,460 white British participants, and an independent GWAS in 42,274 individuals of European ancestry.^13^ The method for estimating SNP effects on vitamin D was described in detail in Manousaki et al.^13^ Briefly, a linear mixed effects model (implemented in the BOLT-LMM software)^26^ was used to estimate the effect of each SNP in the UK Biobank, which was then meta-analyzed with a previously published vitamin D GWAS using inverse variance weighted fixed effect meta-analysis. In both GWAS, 25(OH)D levels were first log-transformed and then standardized. Therefore, all MR effect estimates in this study correspond to 1 standard deviation increase in log-transformed vitamin D level. A total of 138 conditionally independent vitamin D associated SNPs in 69 independent loci were identified by GCTA-COJO v.1.91.1 using the meta-GWAS,^27^ which collectively explained 4.9% of variation in vitamin D and 30% of the total SNP heritability (16.1%).^13^ We used these 138 SNPs in our *primary analysis* as IVs (**Supplementary Table 1)** without further LD clumping to achieve greater statistical power. We additionally performed a sensitivity analysis using the top SNP from each of the 69 independent loci as our *secondary analysis* to ensure strict independence between IVs.

### Data for IV-outcome

We retrieved three collections of GWAS summary statistics from the IEU OpenGWAS Project (https://gwas.mrcieu.ac.uk/): ieu-a and ieu-b, which include GWAS summary statistics generated by various consortia, and ebi-a, which includes GWAS summary statistics imported from the NHGRI-EBI GWAS catalog (https://www.ebi.ac.uk/gwas/). A total of 765 GWAS were included, which represented a comprehensive coverage of human complex traits, diseases and biomarkers. We then filtered these outcome GWAS by retaining those with over 10,000 samples and 1,000,000 SNPs across the genome to ensure statistical power and SNP coverage. Furthermore, we only included the GWAS with the largest sample size if multiple GWAS are available for the same outcome, with the exception of multiple sclerosis (MS); we included four MS GWAS with varying sample sizes and genotyping platforms (genome-wide coverage vs. customized immunochip) to compare the MR causal effect estimates of vitamin D on MS. Finally, we manually curated summary statistics for 26 GWAS that were either not included in the IEU GWAS repository or had a larger sample size compared with the GWAS in the IEU repository for the same outcome. These data filtering and curation strategies yielded a total of 190 eligible outcome GWAS in our MR analysis. All GWAS were conducted in samples of European ancestry, with the exception of two cardiovascular GWAS, which were conducted in mixed populations including both European and Asian individuals. The descriptions, references and sources of these 190 GWAS are listed in **Supplementary Table 2.** These outcome GWAS span a broad phenotypic spectrum, including blood biomarkers; brain imaging measurements; lipid; ophthalmology; cancer; metabolic, cardiovascular, orthopedic, autoimmune, neurological and psychiatric traits and diseases; anthropometric traits; behavioral traits; sleep; cognition; personality; and reproductive traits. For each outcome, we retrieved the variant annotations (SNP rsID, chromosome, position, reference and alternate allele) and summary statistics (effect size, standard error, P-value, allele frequency [if available]) for each SNP instrument using the R package TwoSampleMR (https://mrcieu.github.io/TwoSampleMR/), and harmonized the effects and reference alleles between the vitamin D GWAS and outcome GWAS. For strand ambiguous (palindromic) SNPs, we matched the alleles based on minor allele frequencies (MAF) and discarded those with MAF >0.42 following the default setting in the TwoSampleMR package. The sample sizes of these outcome GWAS ranged from 9,954 (for the manually curated, largest-to-date PTSD GWAS) to 1,030,836 (median 112,561; mean 148,179).

### Statistical analysis

MR uses SNPs as IVs to estimate the causal effect of an exposure on an outcome while controlling for potential confounding, by leveraging the random allocation of alleles at conception. Three assumptions need to be satisfied to ensure a valid IV. The first is the relevance assumption, that is, IVs must be strongly associated with the exposure; the second assumption requires no association between IVs and confounders of the exposure-outcome relationship; and the third is the exclusion restriction assumption, which requires that IVs affect the outcome only through the exposure.^28,29^ If all MR assumptions are satisfied, a causal effect can be estimated based on the observed IV-exposure and IV-outcome associations.

We conducted two-sample MR analyses to test for the causal relationship between circulating 25(OH)D and 190 outcomes. We applied three different MR methods, namely MR-PRESSO,^29^ MR-Egger,^28^ and a weighted median estimator approach,^30^ to estimate causal effects. All three methods are robust to horizontal pleiotropy, i.e., an IV affects the outcome via a separate biological pathway from the exposure under investigation (violation of the exclusion restriction assumption), under the InSIDE (instrument strength independent of direct effect) assumption, and the assumption of at least 50% of the genetic variants being valid instruments (no horizontal pleiotropy). For the robustness of our findings, we considered an estimated causal effect to be significant if at least two out of the three methods showed significant false discovery rate (FDR) corrected P-values. In addition, we performed MR-PRESSO, MR-Egger regression, and modified Q and Q’ tests, to detect bias due to horizontal pleiotropy. MR-PRESSO implements a global test to evaluate the presence of horizontal pleiotropy, and an outlier test to detect specific SNP outliers.^29^ In MR-Egger regression, a significant difference of the intercept from zero suggests the existence of average directional horizontal pleiotropy.^28^ Modified Q and Q’ tests are traditionally used to identify over-dispersion and have been applied in the context of MR to detect outliers caused by horizontal pleiotropy.^31^

### Ethics

Our study is a secondary analysis of existing, de-identified, summary-level GWAS data. Specific ethics for each GWAS examined in this study can be found in the original GWAS publications.

## Results

As shown in **Figures 1–3** and **Supplementary Table 3**, in our primary analysis using 138 conditionally independent 25(OH)D-associated SNPs as IVs, we did not find a causal effect of vitamin D on the outcomes examined here, except for multiple sclerosis (MS) and white blood cell (basophil) count.

**Figure 1.**
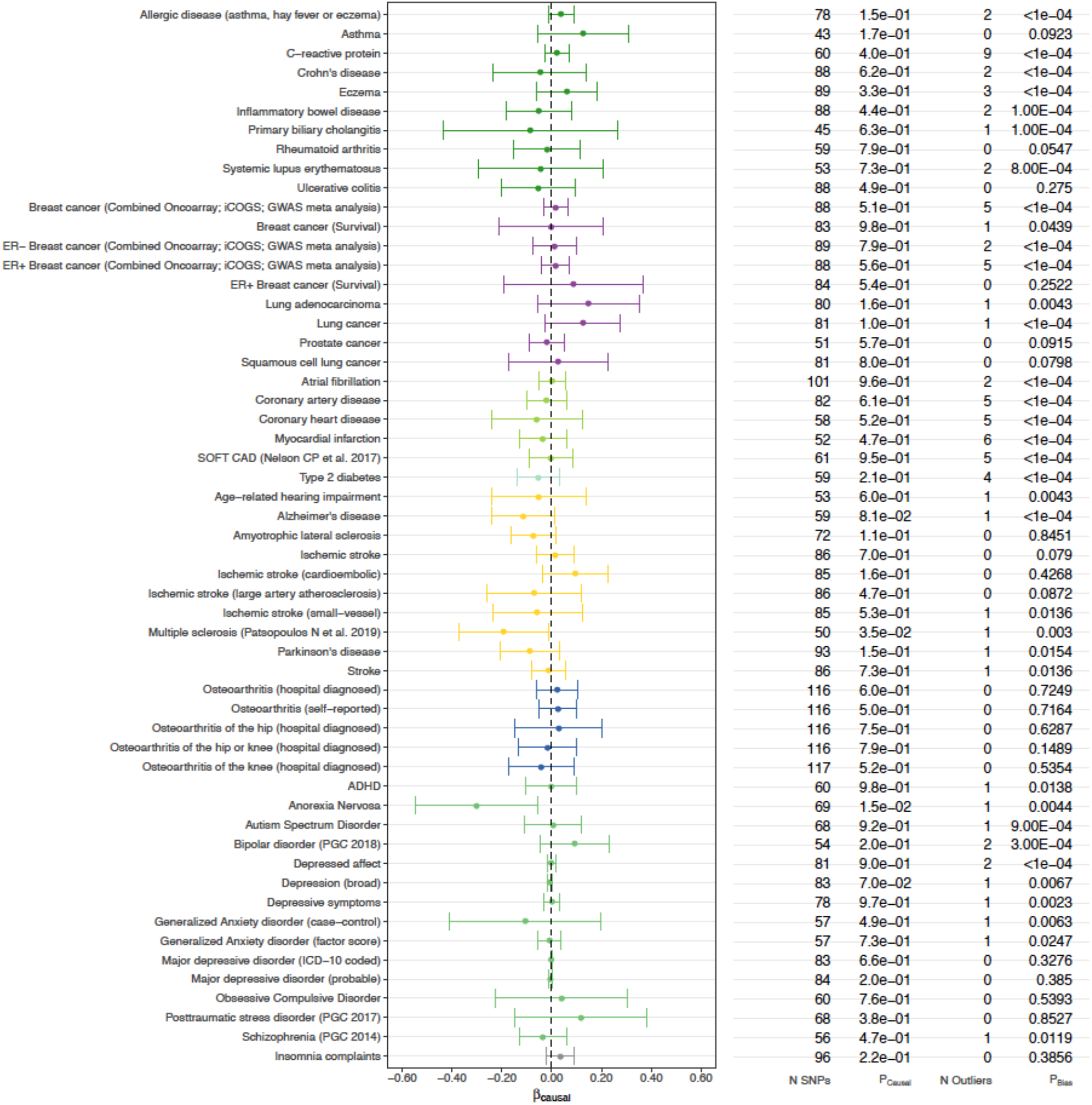
Estimated causal effects of 25(OH)D on *complex diseases* based on the MR-PRESSO approach using 138 genome-wide significant, conditionally independent SNPs of 25(OH)D. The βcausal corresponds to 1 standard deviation increase in log-transformed vitamin 25(OH)D.

**Figure 2.**
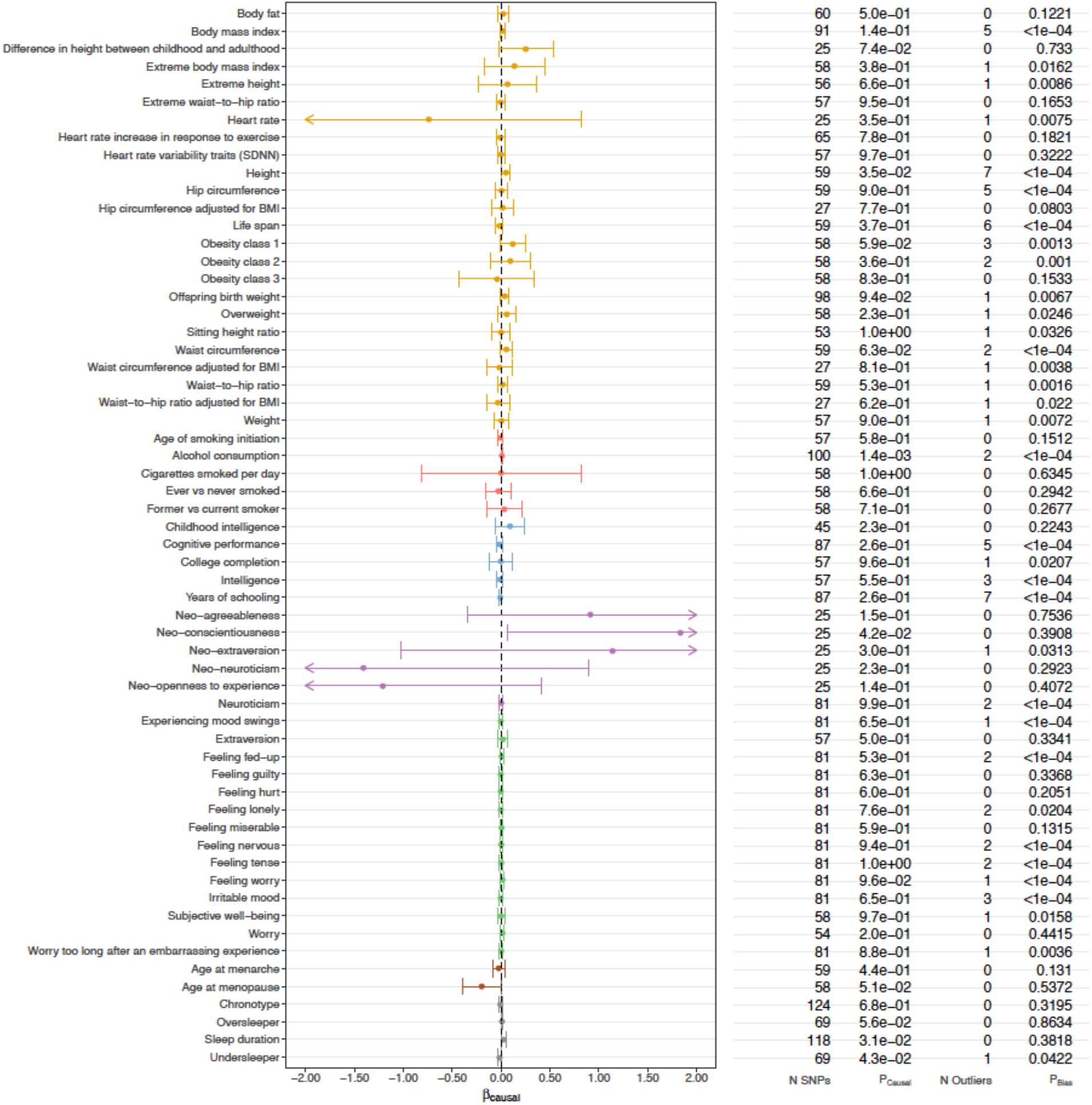
Estimated causal effects of 25(OH)D on *complex traits* based on the MR-PRESSO approach using 138 genome-wide significant, conditionally independent SNPs of 25(OH)D. The βcausal corresponds to 1 standard deviation increase in log-transformed vitamin 25(OH)D.

**Figure 3.**
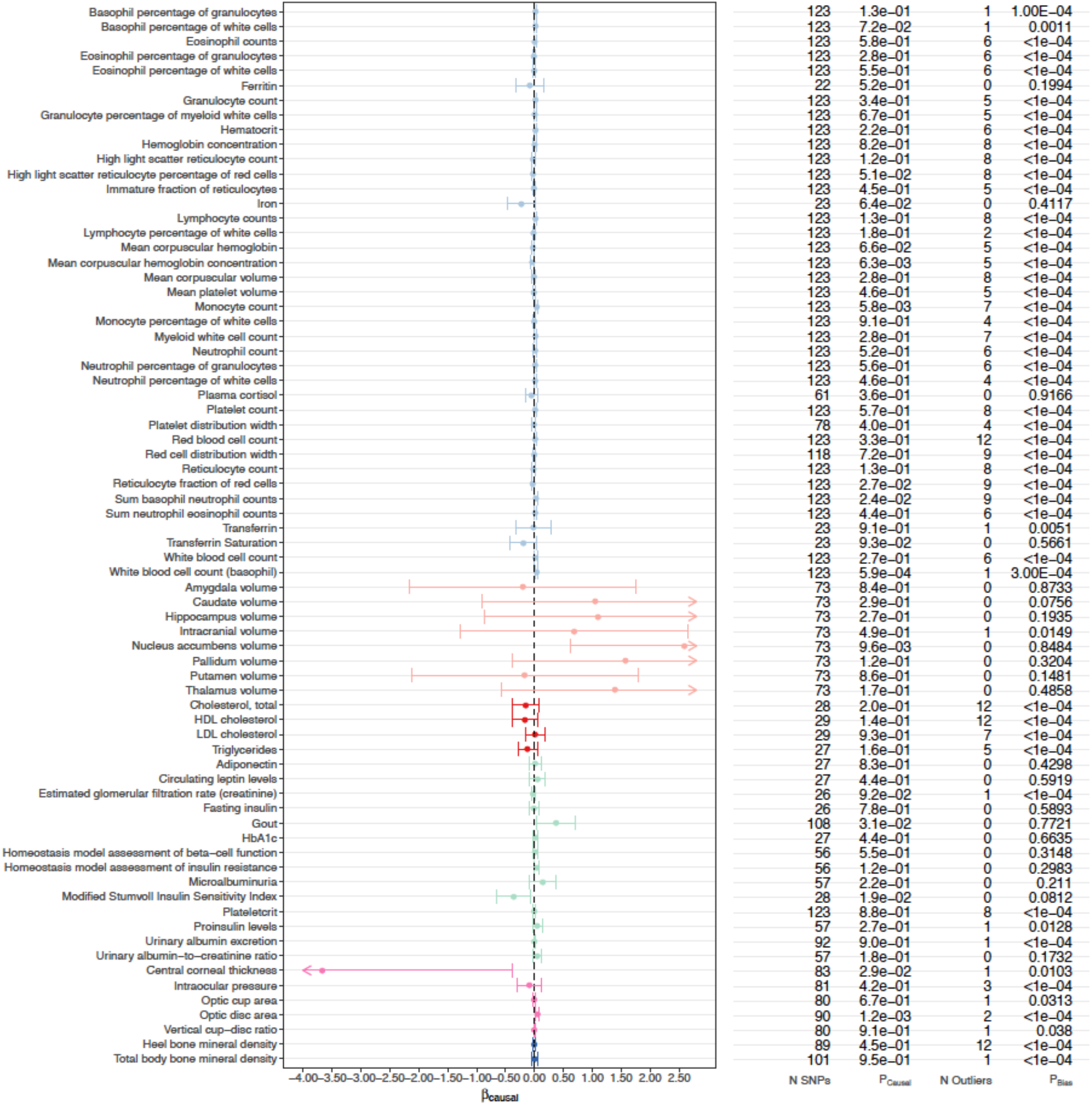
Estimated causal effects of 25(OH)D on *biomarkers* based on the MR-PRESSO approach using 138 genome-wide significant, conditionally independent SNPs of 25(OH)D. The βcausal corresponds to 1 standard deviation increase in log-transformed vitamin 25(OH)D.

Using summary statistics from a study based on Immunochip conducted in 2013, we showed that increased 25(OH)D level decreased the risk of MS (MR-PRESSO odds ratio [OR]=0.433; 95% confidence interval [CI]: 0.313-0.599; P=4.19×10^-7^; P_FDR_=7.96×10^-5^). This result was consistent across different MR methods (MR-Egger: OR=0.390 [0.258-0.588]; weighted median: OR=0.397 [0.301-0.523]) and was in line with previous findings (**Table 1a**). Using an earlier MS GWAS with a smaller sample size (conducted in 2011, 2/3 of the sample size of the 2013 Immunochip study), we observed a similar protective effect, which survived multiple testing correction (MR-PRESSO OR=0.719 [0.606-0.852]; P=1.48×10^-4^; P_FDR_=0.014). We had consistent findings using two other MS GWAS with greater statistical uncertainties (Patsopoulos et al. 2011 GWAS, MR-PRESSO OR=0.705 [0.551-0.900], P=0.005, P_FDR_=0.149; Patsopoulos et al. 2019 GWAS, MR-PRESSO OR=0.824 [0.689-0.986], P=0.035, P_FDR_=0.367) (**Table 1a**). Note that the pleiotropic bias tests suggest that there are various levels of pleiotropic bias for the vitamin D causal effect estimates on MS (**Table 1b**); however, the causal effect estimates reported here are all adjusted for potential pleiotropic bias. The secondary analysis using 69 top SNP instruments also showed consistent results to the primary analysis (**Supplementary Table 4**).

**Table 1.** Causal effect estimates of 25(OH)D and MR pleiotropic bias tests for multiple sclerosis using 138 conditionally independent SNPs in vitamin D associated loci from Manousaki, D. et al. AJHG 2020.

| Outcome | Outcome GWAS sample size | Outcome GWAS N of SNP | N of SNP used in MR | MR-PRESSO |  |  |  | Median estimate |  |  |  | MR-Egger |  |  |  |
| --- | --- | --- | --- | --- | --- | --- | --- | --- | --- | --- | --- | --- | --- | --- | --- |
|  |  |  |  | Beta | 95% CI | P-value | FDR adjusted P-value | Beta | 95% CI | P-value | FDR adjusted P-value | Beta | 95% CI | P-value | FDR adjusted P-value |
| Multiple sclerosis (Patsopoulos N et al. 2011) | 17698 | 7395163 | 66 | -0.350 | (-0.596 - -0.105) | 0.005 | 0.149 | -0.010 | (-0.327 - 0.307) | 0.949 | 0.996 | -0.412 | (-0.742 - -0.083) | 0.014 | 0.226 |
| Multiple sclerosis (Sawcer et al. 2011) | 27148 | 6991436 | 62 | -0.330 | (-0.501 - -0.160) | 1.48E-04 | 0.014 | -0.233 | (-0.429 - -0.036) | 0.021 | 0.436 | -0.422 | (-0.656 - -0.189) | 3.89E-04 | 0.031 |
| Multiple sclerosis (Beecham et al. 2013 immunochip) | 38589 | 1082242 | 24 | -0.837 | (-1.162 - -0.513) | 4.19E-07 | 7.96E-05 | -0.925 | (-1.202 - -0.648) | 6.33E-11 | 1.20E-08 | -0.942 | (-1.353 - -0.531) | 6.90E-06 | 0.001 |
| Multiple sclerosis (Patsopoulos N et al. 2019) | 115803 | 6304359 | 50 | -0.193 | (-0.372 - -0.014) | 0.035 | 0.367 | -0.192 | (-0.365 - -0.019) | 0.030 | 0.436 | -0.234 | (-0.495 - 0.028) | 0.080 | 0.559 |

(b) MR pleiotropic bias tests for 25(OH)D on multiple sclerosis
| Outcome | MR-PRESSO global test P-value | MR-PRESSO N of outlier | Modified Q test P-value | Modified Q' test P-value | MR-Egger intercept test P-value |
| --- | --- | --- | --- | --- | --- |
| Multiple sclerosis (Beecham et al. 2013 immunochip) | 0.029 | 1 | 0.031 | 0.033 | 0.339 |
| Multiple sclerosis (Patsopoulos N et al. 2011) | 0.005 | 1 | 0.006 | 0.005 | 0.575 |
| Multiple sclerosis (Sawcer et al. 2011) | 0.003 | 1 | 0.002 | 0.003 | 0.159 |
| Multiple sclerosis (Patsopoulos N et al. 2019) | 0.003 | 1 | 8.15E-04 | 6.07E-04 | 0.811 |

In addition to MS, we also found white blood cell (basophil) count to be causally affected by vitamin D (MR-PRESSO *β*=0.039, 95% CI: 0.017-0.061, P_FDR_=0.037) (**Figure 3**). However, this result was not significant in our secondary (sensitivity) analysis using 69 top SNPs from independent genomic loci in the vitamin D GWAS (**Supplementary Table 4**). Also note that the pleiotropic bias tests were significant for the vitamin D causal effect estimates on basophil count (**Supplementary Table 4**). Further investigations are thus warranted to elucidate this suggestive finding.

Notably, we identified horizontal pleiotropy (as reflected by the MR-PRESSO global test, the MR-Egger intercept, and modified Q and Q’ tests) for 68% (130 out of 190) of the outcomes analyzed here (123 from MR-PRESSO, 25 from MR-Egger regression, 123 from the modified Q test, and 118 from the modified Q’ test) (**Supplementary Table 3)**.

Lastly, we performed a secondary (sensitivity) analysis using a smaller set of IVs (the top SNP in each of the 69 distinct loci) to ensure the independence between the SNP instruments. As shown in **Supplementary Table 4**, results did not alter substantially by incorporating fewer yet stronger IVs. Most of the outcomes were not influenced by genetically predicted 25(OH)D except for mean corpuscular hemoglobin concentration, which, however, did not survive multiple testing correction in the primary analysis (**Supplementary Table 3 and Figure 3**).

## Discussion

In this study, we performed a comprehensive two-sample MR analysis using 25(OH)D-associated SNPs as genetic instruments and 190 outcome GWAS spanning a wide spectrum of human complex traits and diseases to determine the causal role of 25(OH)D. In line with prior research, we have successfully replicated a causal relationship between 25(OH)D and multiple sclerosis (MS). However, in general, our results, which were based on a markedly increased number of IVs (138 IVs *vs.* 4 or 6 IVs used by most existing MR studies of 25(OH)D) and largely augmented samples of outcome GWAS, showed limited evidence for a causal effect of 25(OH)D on the majority of the traits and diseases investigated here.

The causal role of 25(OH)D on MS has been shown in three independent MR studies, which consistently identified an association between genetically predicted 25(OH)D and a decreased risk of MS.^17–19^ We successfully replicated this causal relationship using a set of updated and more powerful SNP instruments for vitamin D. As a larger number of vitamin D IVs increased the power of the MR analysis, we obtained a more accurate effect size estimate with narrower confidence intervals for the protective effect of 25(OH)D on MS. Although there seemed to be concrete evidence that genetically instrumented 25(OH)D is associated with MS, the causal effects appeared to be decreasing in magnitude as the sample size and SNP coverage increase in the outcome GWAS (50% decreased risk of MS using the 2013 Immunochip study with the smallest sample size; 30% decreased risk using the 2011 GWAS, and 18% decreased risk using the 2019 GWAS with the largest sample size). Noteworthily, our MR finding is based on adult MS GWAS and adult vitamin D GWAS. As a result, the protective effect of vitamin D on MS may only apply to adulthood intervention, although a previous study also suggested a beneficial role of vitamin D for childhood MS.^19^ It is also likely that the protective effect of extra vitamin D on MS is most prominent during critical periods of development (e.g., childhood/adolescence), which cannot be tested by MR analysis based on GWAS using cross-sectional measures of vitamin D in adulthood. Therefore, the causal role of vitamin D in the prevention and intervention of MS needs to be further explored and dissected by future studies.

In addition to MS, a suggestive association was observed for genetically instrumented 25(OH)D and white blood cell (basophil) counts. However, this result needs to be interpreted with caution due to two reasons. First, this effect did not survive multiple testing correction in our secondary analysis using 69 top SNPs from independent loci identified in the vitamin D GWAS. Second, none of the other blood cell counts, including eosinophil, neutrophil, platelet and lymphocyte, appeared to be affected by genetically instrumented vitamin D (all P >0.10). The inverse correlation of vitamin D with the circulating level of basophils has only been reported by one longitudinal cohort study involving 989 young adults.^32^ Despite statistical significance, the correlation coefficient was close to zero indicating a weak effect. Further investigations are thus warranted to confirm this association.

Our null findings are in line with previous MR studies. An earlier MR analysis, which aggregated information from 21 adult cohorts with up to 42,024 participants to explore the causal relationship between vitamin D status and obesity, did not identify any effect of genetically instrumented 25(OH)D on BMI (P=0.57).^33^ In our study, we expanded this outcome category by incorporating several adipose-related phenotypes in addition to BMI, such as body fat percentage, hip circumference, waist circumference, and waist-to-hip ratio (both BMI adjusted and unadjusted). Despite that we used the latest GWAS summary statistics of these outcomes with substantially augmented sample sizes, ranging from 10,255 in extreme waist-to-hip ratio to 315,347 in BMI, we did not find any evidence in support for a causal role of 25(OH)D in obesity-related traits. Likewise, earlier MR studies have reported null findings for most of the autoimmune diseases including inflammatory bowel disease (Crohn’s disease (P=0.67), ulcerative colitis (P=0.42)),^34^ eczema (P=0.27),^35^ lupus (P=0.79) and rheumatoid arthritis (P=0.66).^36^ Our results using summary statistics from greatly enlarged GWAS (for both exposure and outcomes) confirmed a null relationship between 25(OH)D and autoimmune diseases. As for cardiovascular and metabolic traits, previous MR studies did not identify any causal role of 25(OH)D in coronary artery disease,^37^ myocardial infarction,^38^ cholesterol levels including triglycerides and low-density lipoprotein (LDL) cholesterol,^39^ type 2 diabetes,^40,41^ fasting insulin and adiponectin,^42^ all of which, were confirmed by our analyses. There are, however, some discrepancies. For example, an earlier study that incorporated 31,435 individuals and used 4 index SNPs from two 25(OH)D-associated genes found no association between 25(OH)D-lowering alleles and non-fasting remnant cholesterol or LDL cholesterol, but identified a 50% decrease in 25(OH)D levels to be genetically associated with 6.0% (P=0.001) lower high-density lipoprotein (HDL) cholesterol levels.^39^ The effect of 25(OH)D on HDL cholesterol was no longer significant in our analysis when using the summary statistics from a more recent HDL GWAS involving 94,595 individuals (a 3-fold increase in sample size) and a greatly increased number of IVs. Similar null associations were observed in cancers where our MR analysis did not convincingly support a causal effect of circulating 25(OH)D concentrations on the risk of breast or prostate cancer (P=0.58 and 0.46), consistent with previous findings (P=0.47 and 0.99).^43^ Finally, investigations on the causal relationships between 25(OH)D and neurological or psychiatric traits have been limited. So far, only a few MR studies explored the role of 25(OH)D in cognitive function,^44^ major depression,^45^ and Alzheimer’s disease;^20,21^ and nominally significant findings have only been reported in Alzheimer’s disease. It might be worth noting that previous MR studies for Alzheimer’s disease used either IV-exposure association estimates from a smaller 25(OH)D GWAS (SUNLIGHT consortium 2010)^11^ or correlated IVs, which affected the validity of the results. In our updated analysis, the causal effect of 25(OH)D on Alzheimer’s disease attenuated to null (P=0.08 using 2019 Alzheimer’s disease GWAS). Expanding the outcome list by incorporating additional neurological or psychiatric traits also did not identify any significant causal effects. These results are important in informing the ongoing RCT of vitamin D on cognitive change (ClinicalTrials.gov: NCT03613116).

To ensure the validity of MR results, several important assumptions need to be satisfied. First, the relevance assumption, that IVs must be strongly associated with the exposure, is guaranteed by selecting independent 25(OH)D-associated SNPs with genome-wide significance as IVs. The second assumption, that IVs are not associated with confounders of the exposure-outcome relationship, is reassured by the negligible associations between 25(OH)D and various traits and diseases from the current and past MR studies, many of which act as potential confounders for other outcomes (e.g., obesity-related traits). Third, the exclusion restriction assumption requires IVs to affect the outcome only through the exposure (no horizontal pleiotropy). We employed several methods to test and control for horizontal pleiotropy. In particular, MR-Egger regression is designed to detect average pleiotropic bias, whereas the MR-PRESSO global test and modified Q and Q’ tests can detect bias caused by individual outlier SNPs with horizontal pleiotropy. While 68% of the outcomes analyzed here showed significant pleiotropy, we presented MR causal estimates adjusted for pleiotropic bias using three different methods. Notably, the causal effect of vitamin D on MS was also subject to potential pleiotropic bias; however, the effect estimates were significant using multiple different bias-adjusted MR methods.

Our study has several limitations. Sample overlap is a potential concern for our MR analyses, which used genetic instruments derived from the UK Biobank data, given that many recent large-scale outcome GWAS also included UK Biobank samples. Two-sample MR analysis assumes no sample overlap between exposure and outcome GWAS, and the expected covariance of sampling error between the genetic effects on the exposure and the outcome is assumed to be zero. Violating the assumption of no sample overlap in two-sample MR analyses will lead to inflated type I error (false positive) rate for hypothesis testing on the causal effect due to ignoring non-zero covariance of sampling errors.^46^ However, since the majority of our MR results were null, the potentially inflated type I error rate does not change the interpretation of our results. Another limitation of this study is the potential bias in causal effect size estimates due to residual population stratification in the exposure and/or outcome GWAS. However, the vitamin D GWAS we used to extract SNP instruments showed minimal inflation due to population stratification (LD score regression intercept = 1.06; also see Figure S3 in Manousaki et al).^13^ In addition, we leveraged the IEU OpenGWAS Project to extract SNP association statistics from outcome GWAS, which has extensively curated the GWAS summary statistics, with quality control (QC) reports publicly available (as an example, see: https://gwas.mrcieu.ac.uk/files/ieu-a-8/ieu-a-8_report.html). With that being said, we still rely on the quality of the original GWAS for our MR analyses and cannot completely exclude the possibility of biased GWAS association estimates due to insufficient control of population stratification or other suboptimal study designs or analysis strategies, which may bias our MR analysis. Finally, although we leveraged three different MR methods that are robust to pleiotropic bias, all these methods rely on the Instrument Strength Independent of Direct Effect (InSIDE) condition/assumption. Given this limitation, we were not able to completely rule out the possibility that InSIDE condition was violated in some of our analyses, which may result in biased results. However, the three MR methods we used in this study adjust for pleiotropic biases through three different routes, namely pleiotropic bias outliers removal (MR-PRESSO), controlling for average pleiotropic bias (MR-Egger), and robust estimation in the presence of pleiotropic bias outliers (median estimate), and are thus complementary and can provide strong evidence of causal effects when their inferences converge, despite the limitation of the InSIDE condition. Several new methods, such as MR-RAPS and CAUSE, ^47,48^ have been developed, which do not rely on the InSIDE condition. However, these methods require genome-wide GWAS summary statistics, which are not available for some of the exposure and outcome GWAS we examined here, and therefore cannot be applied in a uniform way to investigate the causal effects of vitamin D on the outcomes included in this study.

In summary, our large-scale MR analysis showed limited evidence of the causal effects of vitamin D on a wide range of human complex traits and diseases, except for multiple sclerosis. Our results may inform future RCTs and help to prioritize the most promising target diseases for vitamin D intervention.

## Acknowledgements

We thank the various genomic consortia for generously sharing the genome-wide association summary statistics. This work was supported by the Swedish Research Council [Vetenskapsrådet International Postdoc grant 2015-06522; starting grant 2018-02247], Swedish Research Council for Health, Working life and Welfare [FORTE 2020-00884] (X.J.) and the National Institute on Aging at the National Institutes of Health [grant number K99/R00AG054573] (T.G.). The funders had no role in study design, data collection and analysis, decision to publish, or preparation of the manuscript.

## Author contributions

X.J. and C-Y.C. contributed to study conception, data analysis, interpretation of the results and drafting of the manuscript. T.G. contributed to interpretation of the results and critical revision of the manuscript.

## Competing interests

The authors declare no competing interests.

## Supplementary information

**Supplementary Table 1.** Genetic instruments including 138 conditionally independent SNPs in 69 vitamin D associated loci from Manousaki, D. et al. AJHG 2020 (adopted from Manousaki, D. et al. AJHG 2020)

**Supplementary Table 2.** References for outcome GWAS summary statistics

**Supplementary Table 3.** Causal effect estimates of 25(OH)D on complex traits and diseases and MR pleiotropic bias tests using 138 conditionally independent SNPs in vitamin D associated loci from Manousaki, D. et al. AJHG 2020

**Supplementary Table 4.** Causal effect estimates of 25(OH)D on complex traits and diseases and MR pleiotropic bias tests using top SNPs in 69 vitamin D associated loci from Manousaki, D. et al. AJHG 2020

**Supplementary Table 5.** Causal effect estimates converted to odds ratio for binary traits (138 conditionally independent SNPs)

## Notes

### Competing Interest Statement

The authors have declared no competing interest.

### Summary of Updates

We performed updated MR analyses using new genetic instruments extracted from a meta-analysis combining a vitamin D GWAS conducted in 401,460 white British UK Biobank (UKBB) participants and an independent vitamin D GWAS including 42,274 samples of European ancestry (Manousaki et al. 2020 AJHG) and 190 outcome GWAS. New results showed significant protective effect of vitamin D on multiple sclerosis (MS) across 4 different MS GWAS. However, vitamin D showed no effect on other outcomes examined here.

